# Comparing MEG and EEG measurement set-ups for a brain–computer interface based on selective auditory attention

**DOI:** 10.1101/2024.05.08.593098

**Authors:** Dovilė Kurmanavičiūtė, Hanna Kataja, Lauri Parkkonen

## Abstract

Auditory attention modulates auditory evoked responses to target vs. non-target sounds in electro- and magnetoencephalographic (EEG/MEG) recordings. Employing whole-scalp MEG recordings and offline classification algorithms has been shown to enable high accuracy in tracking the target of auditory attention. Here, we investigated the decrease in accuracy when moving from the whole-scalp MEG to lower channel count EEG recordings and when training the classifier only from the initial part of the recording instead of extracting training samples throughout the recording. To this end, we recorded simultaneous MEG (306 channels) and EEG (64 channels) in 18 healthy volunteers while they were presented with concurrent streams of spoken “Yes”/”No” words and instructed to attend to one of them. We then trained support vector machine classifiers for predicting the target of attention from unaveraged trials of MEG/EEG. Classifiers were trained either on 204 MEG gradiometers or on EEG with 64, 30, 9 or 3 channels and with samples extracted randomly across or only from the beginning of the recording. The highest classification accuracy, 73% on average across the subjects for 1.0-s trials, was obtained with MEG when the training samples were randomly extracted throughout the recording. With EEG, the accuracies were 69%, 69%, 67%, and 63% when using 64, 30, 9, and 3 channels, respectively. When training the classifiers with the same amount of data but extracted only from the beginning of the recording, the accuracy dropped by 12 %-units on average, causing the result from the 3-channel EEG to fall below the chance level. Combining five consecutive trials partially compensated for this drop such that it was 1–8 %-units. While moving from whole-scalp MEG to EEG reduces classification accuracy, a usable auditory-attention-based brain-computer interfaces can be implemented with a small set of optimally-placed EEG channels.

## Introduction

Selective auditory attention modulates the amplitude of the evoked responses to a behaviorally relevant auditory target [1, 2]. Such a modulation recorded non-invasively by magneto-/electroencephalography (MEG/EEG) [1–5] can readily be extracted by machine-learning algorithms and employed e.g. to control a brain-computer interface (BCI) [6–9].

Our previous MEG study [10] utilizing a modified dichotic listening task, where the subjects were presented with two simultaneous streams of spoken-word stimuli, demonstrated that the target of selective auditory attention can be detected at high accuracy from just 1 s of MEG data. However, in this study, as in most MEG/EEG classification studies, the classifier was trained by using data randomly sampled across the entire recording, i.e., the data were *randomly partitioned* for classifier training and testing. Although such a training can be considered optimal, it cannot be applied to a real-time BCI as there the training can happen only with data from the beginning of the measurement session.

Recent studies utilizing auditory EEG-BCI have demonstrated promising results of target detection on healthy subjects [11–16] and on patients [17]. The accuracy of such a BCI naturally depends on the ability of the classifier to extract relevant brain activity [18, 19], which is influenced by the experimental design and measurement set-up [20]: the number of training samples and their distribution across the session, the length of each trial as well as the number, type and location of the measuring channels likely all affect the accuracy but it is not known to which extent.

To explore the effect of these parameters, we recorded simultaneous MEG-EEG while the subjects performed a selective attention task. We investigated how much the classification accuracy decreases when the classifiers are trained only on samples from the beginning of the recording (as in the calibration phase of an actual BCI system) instead of using the same number of samples randomly extracted from the entire recordings (as in typical offline classification). Moreover, we tested how the classification accuracy depends on the type (MEG vs. EEG) and number of channels and on the number of trials used for training the classifiers.

## Materials and methods

### Participants

Eighteen healthy adult volunteers (8 females; 10 males; mean age 28.8 ±3.8 years, range 23–38 years) participated in the study. According to Edinburgh Handedness Inventory [21], one subject was left-handed, three were ambidextrous and the rest were right-handed. Subjects did not report hearing problems or history of psychiatric disorders. The study (statement “2017-15-Real-time detection and decoding of responses”) was approved by the Aalto University Ethics Committee. The research was carried out in accordance with the guidelines of the Declaration of Helsinki, and the subjects gave written informed consent prior the measurements. One subject was excluded from MEG/EEG data analysis due to misunderstanding the task, one was excluded from MEG data analysis due to poor data and six more were excluded from EEG data analysis due to extensive artefacts in their data leaving sixteen test subjects for MEG and eleven test subjects to EEG. When comparing MEG and EEG, only data of the same subjects were contrasted.

### Stimuli and experimental protocol

The stimuli were adapted from our previous study [10]. The stimuli created an acoustically realistic scene of two speakers standing in front of the subject at –40 (left) and +40 degrees (right) from the midline, continuously uttering the words “Yes” (a female speaker) and “No” (a male speaker) in a sequence where high- and low-pitch versions of the stimulus word alternated. Both words were presented once per second, and the streams were interleaved such that the words had minimal overlap; see Fig. 1. The word sequences contained occasional deviants (violations of the regular alternation) that comprised three consecutive high-pitch versions of the word and occurred with a 5% probability in each stream; see Fig. 1c.

**Fig 1.**
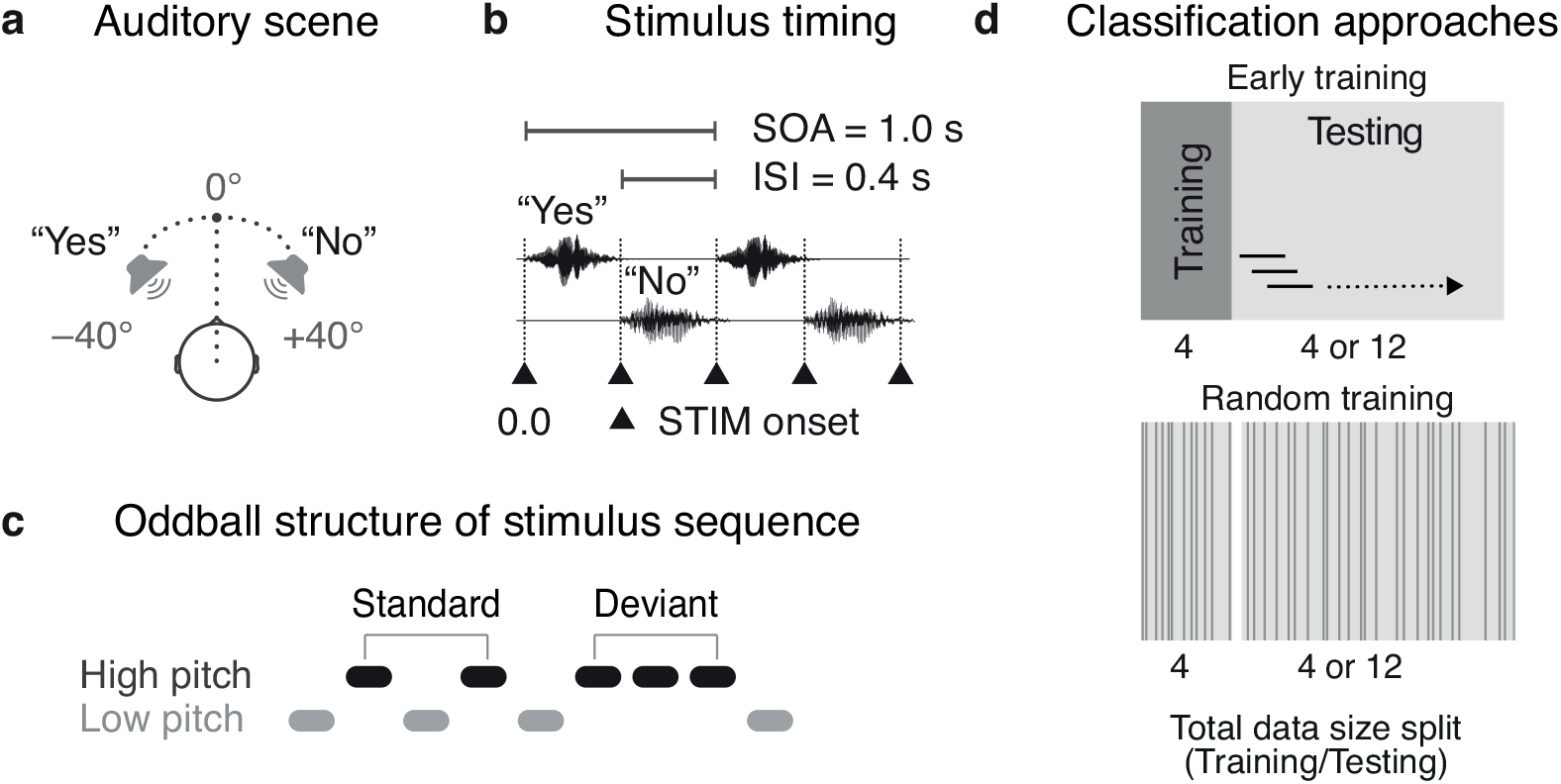
Experimental design and conditions. A: Spoken-word stimuli recorded with a dummy head were presented to the subject. In the acoustic scene, the two speakers appeared at +/– 40 degrees of the subject’s head mid-line. B: The stimulus onset asynchrony (SOA) was 1 s, leaving a 400-ms interval (ISI) between consecutive stimuli. The black triangles indicate stimulus onsets. C: The high- and low-pitch version of the words alternated (standard) but occasionally two same-pitch words were presented consecutively (deviant). D: Classifiers were trained with samples either only from the beginning of the recording (early training) or with samples extracted randomly across the entire recording (random training).

PsychoPy (version 1.79.01) [22, 23] Python package was used to control the experiment and to present the auditory stimuli and visual instructions. PsychoPy was run on Ubuntu Linux (version 14.04). Auditory stimuli were delivered by a professional audio card (E-MU 1616m PCIe; E-MU Systems, Scotts Valley, CA, USA), an audio power amplifier (LTO MACRO 830; Sekaku Electron Industry Co., Ltd, Taichung, Taiwan), custom-built loudspeaker units outside of the shielded room, and plastic tubes conveying the stimuli to both ears separately. The sound pressure was adjusted to a comfortable level for each subject individually.

### Experimental structure and the task

The experiment comprised 16 blocks and in total lasted for 45 minutes including the breaks between the blocks.

The task of the subject was to focus on the spoken-word stream indicated in the visually-presented instruction prior to each block and to minimize eye movements by maintaining gaze at the fixation cross displayed on the screen, which was 1.4 m from the eyes of the subject. If the cue “LEFT-YES” was displayed, subjects had to attend to the “Yes” stream presented by the virtual speaker on their left side, and conversely for the cue “RIGHT-NO”. The experiment always started with the block “LEFT-YES”, followed by the block “RIGHT-NO”. The subsequent blocks were presented in a random order for every subject.

### MEG/EEG data acquisition

MEG/EEG measurements were performed with a whole-scalp 306-channel Elekta–Neuromag VectorView MEG system (MEGIN Oy, Helsinki, Finland) at the MEG Core of Aalto Neuroimaging, Aalto University. All subjects were measured in the seated position. In addition to MEG, simultaneous EEG was recorded with a 64-electrode Waveguard™ EEG cap (Advanced NeuroTechnology, Enschede, The Netherlands).

Anatomical landmarks (nasion, left and right preauricular points), head-position indicator coils, and additional scalp-surface points (around 100) were digitized using an Isotrak 3D digitizer (Polhemus Navigational Sciences, Colchester, VT, USA). Bipolar electrooculogram (EOG) with electrodes positioned around the right eye (laterally and below) was recorded. MEG/EEG data were filtered to 0.03–330 Hz and sampled at 1 kHz during recording.

### Analysis

The analysis pipeline was implemented with MNE-Python (version 0.17) [24] and scikit-learn (version 0.18) [25] software packages.

### Pre-processing

External magnetic interference in the MEG signals was suppressed using signal-space separation [26] as implemented in the MaxFilter software (version 2.2.10; MEGIN Oy, Helsinki, Finland). No continuous head movement correction was performed because the MEG data were recorded without continuous head position tracking. The EEG data were transformed to average reference after removing bad channels. Bad channels were not interpolated because averaging the neighboring activity to recover the bad channel(s) would not add additional information for the classifiers. The number of bad channels varied from 2 to 11 (only single subject), on average 4.3 channels, which leaves on average 60 good channels for EEG analysis, however in the text EEG will be always referred as 64-channel EEG.

The unaveraged MEG-EEG data were down-sampled to 125 Hz and filtered to 0.1–45 Hz. Ocular and cardiac artefacts were suppressed by removing those independent components (3–10 per subject, on average 6) that correlated most with the EOG or ECG signals, respectively. One-second epochs (–0.2, 0.8 s) were extracted from the MEG/EEG data around every stimulus and baseline-corrected. Individual epochs were discarded if their signal amplitude exceeded 5 pT for magnetometers, 4 pT*/*cm for gradiometers, 80 *µ*V for EEG and 150 *µ*V for EOG. Epochs corresponding to deviants were excluded from further analysis.

After pre-processing, one measurement block yielded on average 356 accepted trials (range 268–422 trials).

### Classification

Prior to passing the MEG/EEG signals to the classifier, their mean was removed and they were scaled to unit variance.

Two linear support vector machines (SVM) [27] were applied on the down-sampled epochs for the classification task. One SVM was used for classifying attended vs. unattended stimuli in the “Yes” spoken-word stream and similarly another one for the “No” stream.

We trained the SVM classifiers in two different ways. First, similarly to the calibration phase of an actual BCI, we trained the classifiers with the data samples extracted only from the beginning of the measurement (corresponding to the first four data blocks B1–B4), later referred to as *early training* ; see Fig. 1d. Second, we trained the classifiers by randomly picking the training data samples over whole measurement, later referred to as *random training*; see Fig. 1d. In both training modes, 25% of the data, which is equal to four data blocks, were used for the training.

The early-training classifier utilized test data from blocks B5–B16 in three different ways. In the first way, the classification results for five consecutive trials were combined to yield a confidence value ranging from 0 (full confidence for the “Yes”-word stimuli) to 1 (full confidence for the “No” stream). The value of 0.5 indicated that the classifier had no information on the direction of auditory attention. This combination was done as a moving average spanning 5 trials (5 s of data) at a time. The second way was like the first one except only two consecutive trials were combined. Finally, the third way was to employ only single trials.

The effect of varying the number of EEG channels used for the classification was investigated by restricting the original 64-channel EEG to 30 channels (electrode locations according to the international 10–20 system), to nine channels (*Fz, FC1, FC2, Cz, C3, C4, CP1, CP2, Pz*) and to three channels (*Fz, Cz, Pz*). The nine- and three-channel selections were determined based on a study by Sellers and colleagues [28]. The different EEG channel sets are shown in Fig. 2.

**Fig 2.**
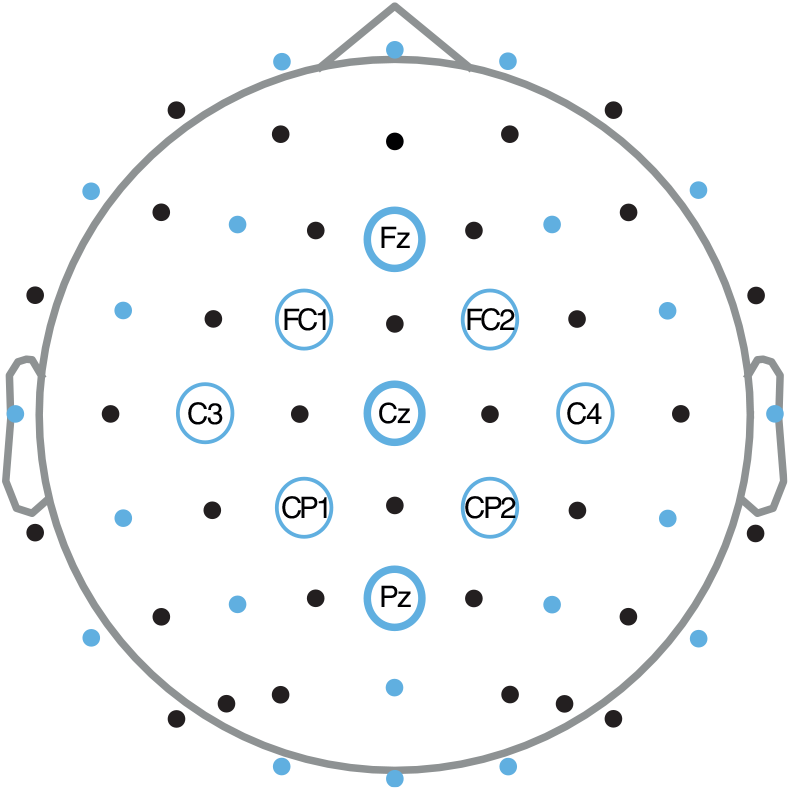
Layout of the 64 EEG electrodes in the measurement (all dots), and the subsets of 30 (blue dots and all circles), 9 (all blue circles) and 3 (thick blue circles) channels.

The dependence of classification accuracy on the amount of training data was tested by varying the number of training samples with p-left-out single-SVM classifier. The classification training performed for training sizes starting with 10 samples and gradually increasing it with a step of 50 samples to the maximum of training data samples (750 samples) with the 20 iterations in each step. The data before splitting into training (750 samples) and test (250 samples) data sets were randomly shuffled. Then training samples for each step (from 10 samples up to 750 samples) were selected randomly from the training data set and repeated in each iteration in range of twenty. At each step, the classifier was tested on 250 left-out samples. The average classification accuracy and standard deviation were calculated for each step across those 20 iterations. The classifiers were tested on 204-channel MEG and 64-, 30-, nine-, and three-channel EEG selections.

## Results

MEG yielded the highest classification accuracy, reaching 73% for single (1.0-s) trials when training with randomly extracted samples; see Fig. 3. In the same conditions, the 64-channel EEG provided the slightly lower accuracy of 69.0%, which decreased only little when reducing the number of channels to 30 (68.6%) and 9 (66.8%) but dropped more substantially with just 3 channels (61.1%).

**Fig 3.**
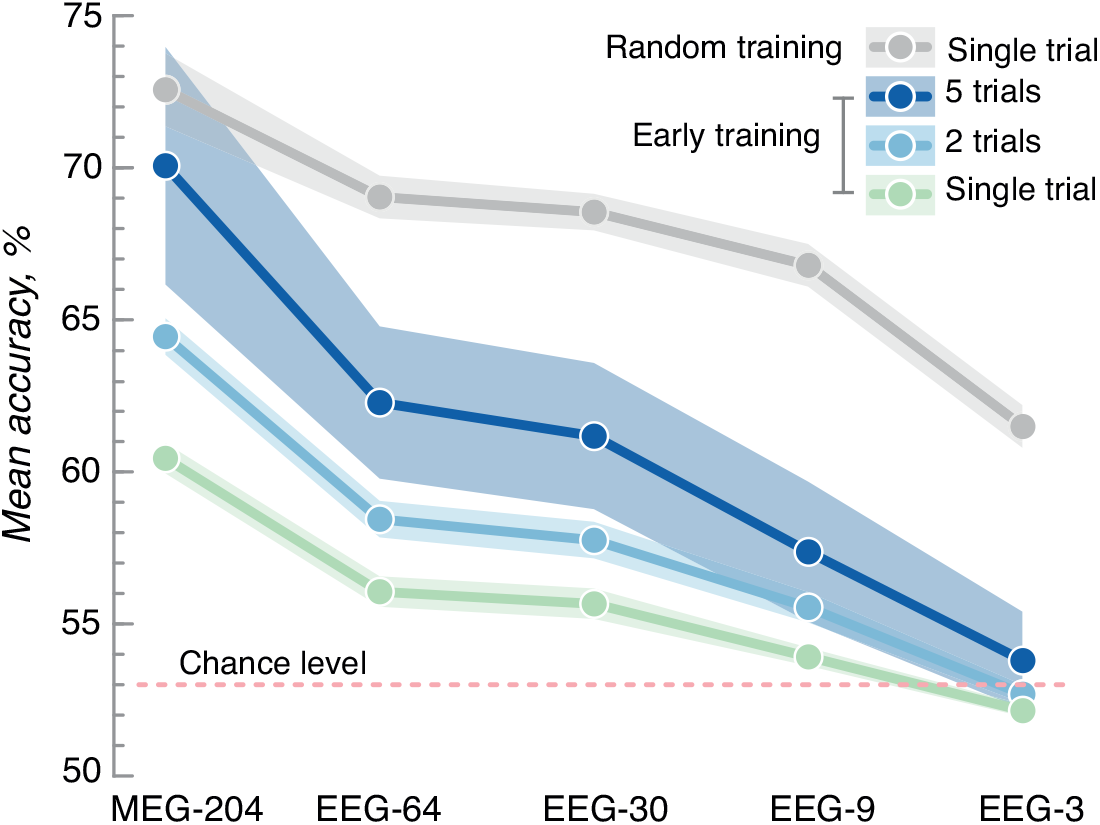
Classification accuracy as a function of the measurement set-up and the number of combined trials. Random training (grey) refers to extracting the training samples randomly across the entire recording while in early training (blue, green) the classifiers were trained on the first quarter of the data. The dots represents the mean accuracy across the 11 subjects, and the shading the standard error of mean.

Classifiers trained with samples extracted randomly across the recording (as is customary in offline classification studies) yielded a systematically higher accuracy than those trained with the same number of samples only from the beginning of the recording (as must be done in an online BCI application); see Fig. 3. This difference was on average 12 %-units (range 9–13 %-units) across the measurement set-ups.

Combining the classification results of consecutive single trials improved the overall classification accuracy as anticipated. Here, by using two trials increased the accuracy by around 2 %-units on average across the measurement set-ups and by using five trials by around 5 %-units.

To find out the required number of training trials for a given decoding accuracy, we trained a classifier with a variable number of trials and tested with 250 left-out samples. This test-set accuracy is shown in Fig. 4. For example, to achieve 70% classification accuracy on 204-channel MEG, about 130 training trials (about 2 min) were needed. To reach the same accuracy on 64-, 30-, and 9-channel EEG required about 310 trials (about 5 min). With only 3 EEG channels, over 510 trials (9 min) were required for classifier training to reach that 70% accuracy.

**Fig 4.**
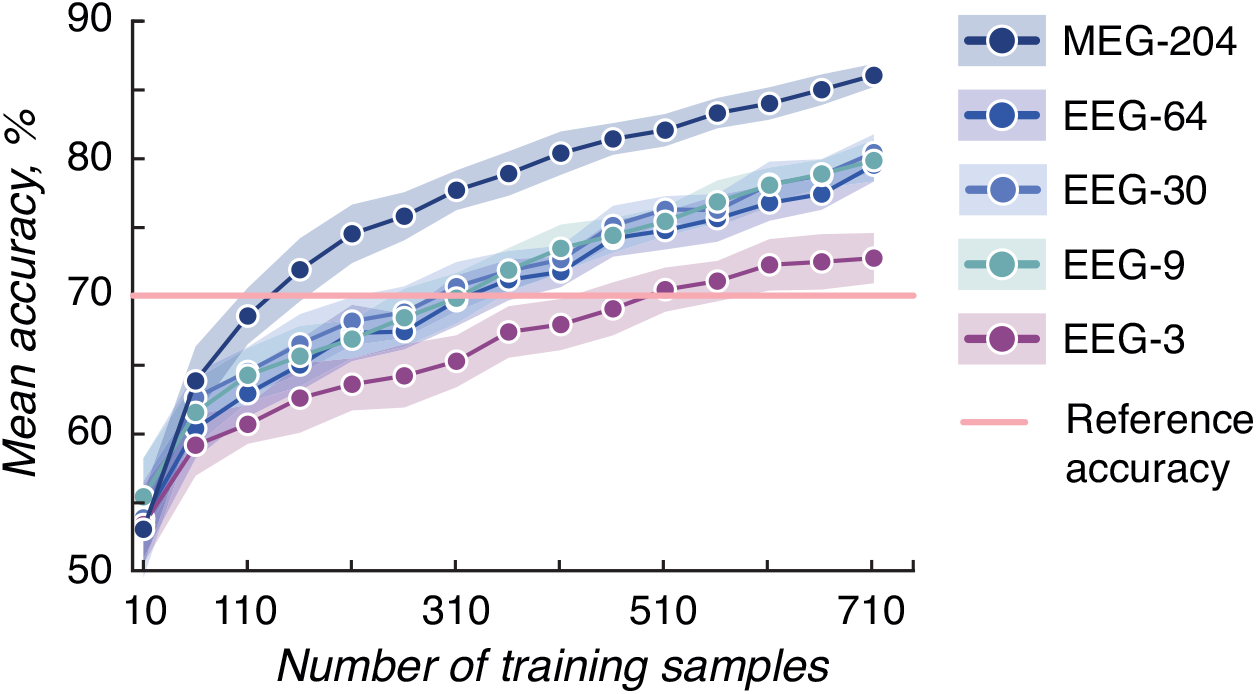
Classification accuracy of the direction of auditory attention as a function of the amount of training data. The classification learning curves are shown for a representative subject (S6). The shading indicates the standard deviation calculated over 20 iterations to each training sample size.

## Discussion

In this study, we investigated how the performance of an auditory-attention-based BCI depends on the measurement modality (MEG vs EEG), the number of measurement channels (EEG), the number of trials combined for one classification result, the number of training samples and whether they are extracted randomly across or only from the beginning of the recording as must be done in an actual BCI.

MEG yielded a significantly better classification accuracy than the EEG set-ups we included in this study. Since we utilized 204 MEG channels and at maximum only 64 EEG channels, this comparison does not directly tell about the relative performance of MEG vs. EEG. However, since increasing the EEG channel count from 30 to 64 led only to a small increase in the classification accuracy, we would anticipate that increasing the EEG channel count further to 128 or even higher would not make EEG to outperform MEG. Yet, this performance advantage of MEG over EEG is mostly theoretical, since MEG is costly and non-portable and thus not well suited for BCIs outside of the laboratory environment.

As expected, reducing the number of EEG channels utilized in the classification task impacted the classification accuracy considerably when the channel count was low to begin with. Our results with a 9-channel EEG did not indicate sufficient performance for a practical BCI even though the locations of these channels were optimized for this task based on previous studies. Yet, surprisingly, the offline classification (training samples extracted randomly across the recording) yielded acceptable performance with just those 9 channels. Classifying attention direction from the 3-channel EEG did not exceed the chance level even when combining two trials.

The classification accuracies obtained with the 64- and 30-channel EEG were rather similar, indicating that a 30-channel system — even without task-optimized electrode locations — is already able to sample almost all of the information available in scalp EEG about the direction of auditory attention. These results are in line with previous study examining the optimal number of EEG channels for comparable classification tasks; they have demonstrated that the marginal utility of additional channels drops above about 22 channels [19].

Cross-validation-type training and testing of the classifier gave substantially higher accuracy than training with the same number of samples exclusively from the beginning of the recording. This difference could be due to the participants changing their strategy of attending one stimulus stream in the course of the experiment. However, it could also stem from within-measurement changes in background brain activity, physiological artifacts and ambient interference, which would all make training the classifier based only on the beginning of the measurement suboptimal. A likely neurophysiological change is the increase of on-going alpha- and beta-band activity as the subject gets more relaxed or even drowsy as the measurement proceeds and the initial excitement is over.

For a random-training classifier to reach 70% accuracy, almost 2.4 times more training trials of 30- and 9-channel EEG was required compared to 204-channel MEG. This difference is likely mostly due to the mere difference in the number of measurement channels but possibly also due to the more confined and direction-dependent sensitivity patterns of the planar gradiometer channels of MEG vs EEG channels.

The loss of classification accuracy due to fewer measurement channels or shorter training time can be partly compensated for by combining classification results of consecutive trials at the expense of the information transfer rate (ITR) such a BCI can provide. However, combining results of individual trials increases the overall classification accuracy only if there is no systematic classification error. Our results speak for no considerable bias in the classification results of individual trials since we observed a clear and monotonic increase in accuracy when adding more trials to the combination. To optimize the trade-off between ITR and classification accuracy, da Cruz and colleagues [29] proposed an adaptive decision time window, which improved ITR by 19% in their BCI based on steady-state visual evoked potentials. In our experimental design, an adaptive time window could be realized by accumulating classification results from as many trials as needed for reaching a certain confidence level, determined based on the class probabilities given by the SVM algorithm.

In this study, we did not aim at a BCI that would generalize across participants but instead trained the BCI individually for each participant. Such individual training has been shown to yield higher performance also in certain patient populations [30]. However, the ability of the BCI to generalize across subjects could still be a useful feature [31, 32] since an individually-tuned BCI takes time calibrate, however, it typically provides a higher classification accuracy than a generalizing BCI.

We did not focus nor evaluate factors such as motivation, mental fatigue, frustration and anxiety [33–36] or the mental strategy participants employed to focus attention. These factors may have an effect on classification accuracy [36].

## Conclusion

In conclusion, we have demonstrated how training time and moving from whole-scalp MEG to EEG with different channel counts affects classification accuracy in an auditory-attention-based BCI. Our results provide guidelines on how to reach the desired classification accuracy given the measurement set-up.

## Acknowledgements

This research was supported by Academy of Finland, grant No. 295075 “NeuroFeed”, and European Research Council, grant No. 678578 “HRMEG”. The content is solely the responsibility of the authors and does not necessarily represent the official views of the funding organizations. The measurements were conducted at the MEG Core of Aalto Neuroimaging Infrastructure, Aalto University, Finland, and financially supported by Aalto Brain Centre. The authors thank prof. Ville Pulkki and Aalto Acoustics Lab, Aalto University, for the loan and guidance on the use of the dummy head.

